# Engineered Active Zymogen of Microbial Transglutaminase

**DOI:** 10.1101/2023.10.09.561484

**Authors:** Ryutaro Ariyoshi, Takashi Matsuzaki, Ryo Sato, Kosuke Minamihata, Kounosuke Hayashi, Rie Wakabayashi, Masahiro Goto, Noriho Kamiya

## Abstract

Microbial transglutaminase (MTG) has shown to be a powerful biocatalytic glue for site-specific crosslinking of a range of biomolecules and synthetic molecules, those handled with an MTG-reactive moiety. The preparation of active recombinant MTG requires the posttranslational proteolytic digestion of propeptide working as an intramolecular chaperon to assist the correct folding of MTG zymogen (MTGz) in the biosynthesis. Herein, we propose an <u>e</u>ngineered active <u>z</u>ymogen of MTG (EzMTG) that is expressed as soluble form in the host *E. coli* cytosol and exhibits the cross-linking activity without limited proteolysis. Based on the 3D structure of MTGz and serendipitous findings, saturated mutagenesis of K10 or Y12 in propeptide domain leads to generate several active MTGz mutants. In particular, K10D/Y12G mutant exhibited the catalytic activity comparable with a mature form. However, the expression level was low possibly due to the reduction of chaperone activity and/or the promiscuous substrate specificity of MTG, which is potentially harmful to the host cells. By contrast, soluble K10R/Y12A mutant is expressed in the cytosol of host *E. coli* and exhibited unique substrate-dependent reactivity toward peptidyl substrates. The quantitative analysis of the binding affinity of mutated propeptide to the active site suggested the trade-off relationship of EzMTGs between the binding affinity and the catalytic activity. Our proof-of-concept study provides insights into the design of a new biocatalyst by using the zymogen as a scaffold and will convey a potential route to the high-throughput screening of MTG mutants for bioconjugation applications.

## Introduction

Enzymes are natural proteinaceous catalysts showing diverse reaction specificity and thus are widely utilized from basic science to practical applications. Biosynthesis of enzymes is often linked to the activity regulation in the host organisms. The control of enzymatic activity is vital for the spatio-temporal control of both activation and inhibition to sustain the cellular physiology. Specifically, proteases are often installed as a precursor of the mature form (zymogen) to control the proteolytic digestion of proteinaceous endogenous constituents in the host organisms.^1,2^ Nature adapts several ways to regulate proteolysis intermolecularly by using endogenous proteinaceous inhibitor or intramolecularly with propeptide that works as an intrinsic inhibitor and supports the correct folding. In the latter case, limited proteolysis of the propeptide of zymogen generates an active, mature enzyme. Interestingly, zymogen could show weak reactivity to a specific substrate^3^, implying the modulation of interaction of propeptide with the enzyme active site may increase the frequency of substrate entry to the active site. In fact, enzymatic activity of some protease was markedly enhanced by the deletion of several amino acids adjacent to propeptide^3^. The accumulated knowledge on the natural and artificial strategies to modulate activity of zymogen motivated us to engineer the scaffold of zymogen as a base structure for the design of a new biocatalyst.

Transglutaminase (TGase, protein-glutamine γ-glutamyltransferase, EC 2.3.2.13,) widely found in natural organisms, such as microorganism^4^, plants^5^, insects^6^, and mammals^[7-8]^, works as a natural glue by crosslinking the γ-carboxyamide groups of glutamine (Q) and a variety of primary amines, including the ε-amino group of lysine (K) ^4,9^. Microbial transglutaminase (MTG) from *Streptomyces mobaraensis* is a stable, Ca^2+^-independent enzyme^10–12^. Owing to the promiscuous specificity for proteinaceous substrates, MTG has rapidly distributed in food industry as edible glue^12^. MTG has also been recognized as a powerful tool in bioconjugation applications, such as crosslinking of single-chain antibody and enzyme^[13]^, bioconjugation of protein of interest (POI) with small chemical entities^14–17^ or with synthetic macromolecules such as polymers bearing MTG-reactive moieties.^18–20^ and chemically modified nucleic acids.^21^. Recently, MTG has been actively employed in the preparation of antibody-drug conjugates (ADCs) and bioimaging ^22–24^, envisaging further biological and biotechnological applications in biopharmaceutical industries.

The promiscuity in the substrate recognition of MTG, however, limits its utility as a versatile biocatalytic glue especially for *in vivo* applications. MTG expresses as an inactive zymogen with the propeptide that functions intramolecular chaperone which assists correct folding ^25–28^ as well as shields the host cells from its deleterious crosslinking of endogenous proteins. The expression of active MTG by microorganisms is thus associated with limited proteolysis upon the secretion^29,30^. In the case of *E. coli*-based expression system, *in vitro*^31^ or *in vivo*^32,33^ posttranslational treatment by a specific protease is required to obtain active recombinant MTG. Alternatively, co-expression of genes encoding propeptide and mature MTG was found to yield the active form, though additional purification steps are needed to separate the two components.

To dissect the substrate specificity of MTG, researchers explored highly reactive peptidyl substrates by using a phage display system^34^ and designed a pair of engineered small proteinaceous substrates and MTG mutants by protein engineering.^35^ Although combination of rational design and screening should be an important option for further engineering of MTG, difficulty in the expression of active mature MTG mutant in the cytosol of host cells such as *Escherichia coli* ^33^ hampers the feasible construction of mutant libraries to tune its substrate specificity.^36^ The high-throughput screening (HTS) of active MTG by posttranslational proteolysis of zymogen mutants displayed on yeast surface was investigated,^37^ yet still difficult to validate the substrate specificity of the soluble MTG mutants in an HTS manner. In this context, another glue enzymes with narrower substrate specificity in comparison with MTG such as sortase A have widened the utility for researchers^[38]^. Searching other organisms producing TGase potentially applicable to highly specific bioconjugation is also an important option and KalbTG isolated from *Kutzneria albida* was reported to be compatible to the preparation of ADCs than MTG.^39^

Herein, we propose a novel <u>e</u>ngineered active <u>z</u>ymogen of <u>MTG</u> (EzMTG) (**Fig. 1**). Our concept to create an active, soluble recombinant MTG that can be expressed in *E. coli* cytosol is to simply substitute two amino acids, K10 and Y12, in the propeptide to relieve the interaction with the active site of MTG for the substrate entry while keeping its intramolecular chaperone functionality. The saturation mutagenesis of each amino acid leads to the design of K10D/Y12G mutant that exhibits the crosslinking activity comparable to that of the mature counterpart. We also found that K10R/Y12A mutant exhibits higher reactivity toward peptidyl substrates than small molecular substrates, by which the possibility of substrate-dependent switch-on crosslinking activity was shown. Our results will open one way to generate new MTG mutants with altered substrate specificity, which would illustrate the potential of engineering the scaffold of naturally occurring enzyme precursors toward new enzymatic function.

**Fig. 1.**
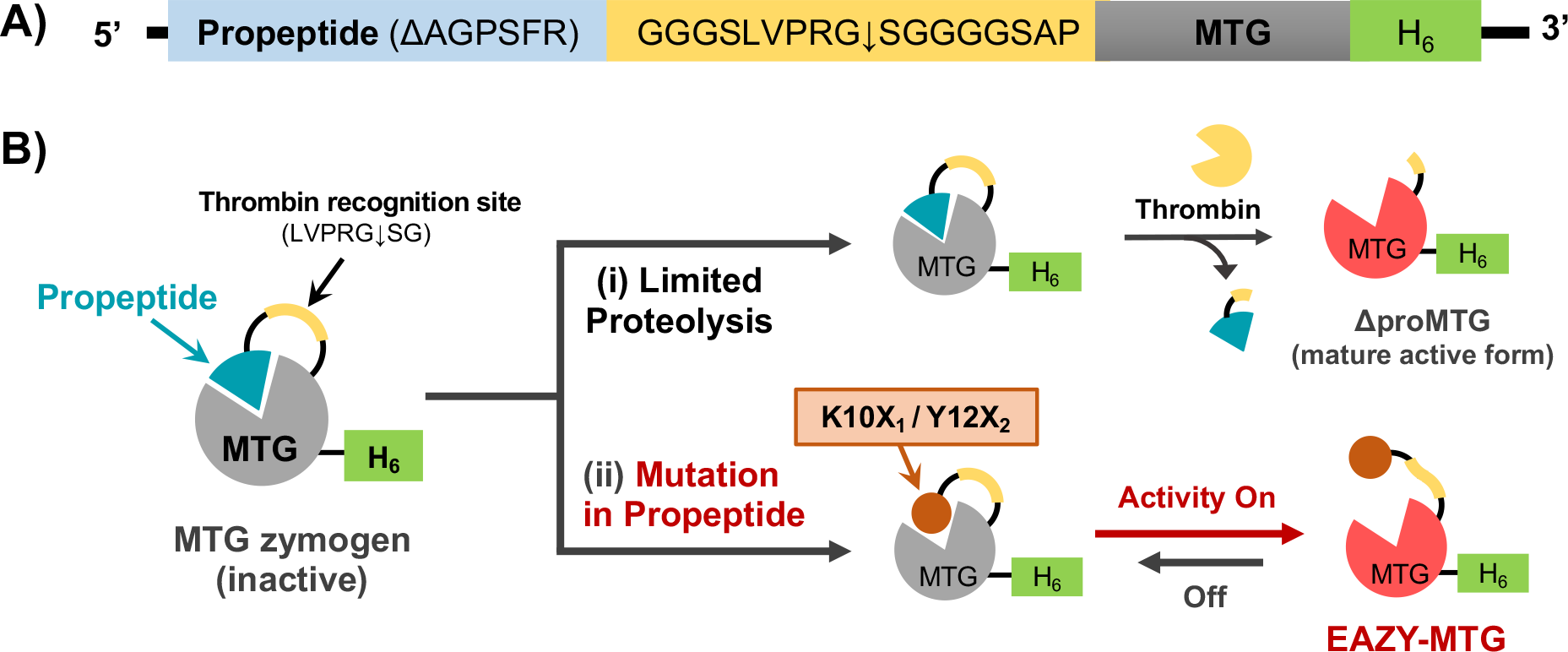
Constructs of EzMTG evaluated in this study. (A) Schematic illustration of DNA constructs of EzMTG. (B) Schematic illustration of the preparation of recombinant active mutants from MTG zymogen (MTGz) (i) by limited proteolysis by thrombin or (ii) designing mutated propeptide to modulate the interaction with the active site.

## Results and discussion

### Design of EzMTG based on the zymogen scaffold

To validate our concept shown in **Fig. 1**, a series of mutants of recombinant MTG zymogen were designed (**Fig. S1**). First, we designed the base protein scaffold of EzMTG by replacing a part of C-terminal sequence (AGPSFR) of the propeptide domain with a thrombin-digestive site encompassed with a Gly-Ser flexible linker (GGGS LVPRGS GGGS). The insertion of the flexible linker is expected to facilitate the proteolysis by thrombin to obtain a recombinant mature MTG with C-terminal His_6_-tag (ΔproMTG). ΔproMTG is used as the positive control in this study. Next, we selected Y12 as the starting point to engineer the scaffold of zymogen by referring the results of crystal structure analysis of MTG precursor, showing that Y12 is crucial for strong binding between the propeptide and the active site of the mature form.^40^ Unexpectedly, we found that the single mutation of Y12 to Ala (Y12A) makes MTG self-reactive as this mutant was self-labeled with a MTG-reactive fluorescent peptidyl substrate, FITC-β-Ala-QG (**Fig. S2**). The results imply that the single mutation (Y12A) could promote the activation of MTG zymogen. Further studies showed that the replacement of K10 totally suppressed the self-labeling (**Fig. S3**). On the basis of the serendipitous findings, two amino acids, K10 and Y12, located in the propeptide domain were selected for the mutation sites to design EzMTG.

### Expression and enzymatic activity of K10X/Y12A mutants

We first prepared K10X/Y12A MTGz mutants by saturation mutagenesis of K10 to 19 natural amino acids. The initial screening of catalytic activity was done by a standard hydroxamate assay using Z-QG and hydroxylamine (**Fig. S4 (i)**). **Figure 2** showed that the replacement of K10 with D or E increased the catalytic ability, implying the presence of electrostatic repulsion around the K10 position on the protein surface, thereby facilitating the recognition of small substrates. K10G/Y12A mutant also showed comparable activity with K10E/Y12A mutant. By contrast, the low catalytic activity of K10R/Y12A and K10H/Y12A mutants suggests that mutated propeptides remained native-like structure by the substitution of K10 with basic amino acids. Other 14 mutants showed comparable low catalytic activities.

**Fig. 2.**
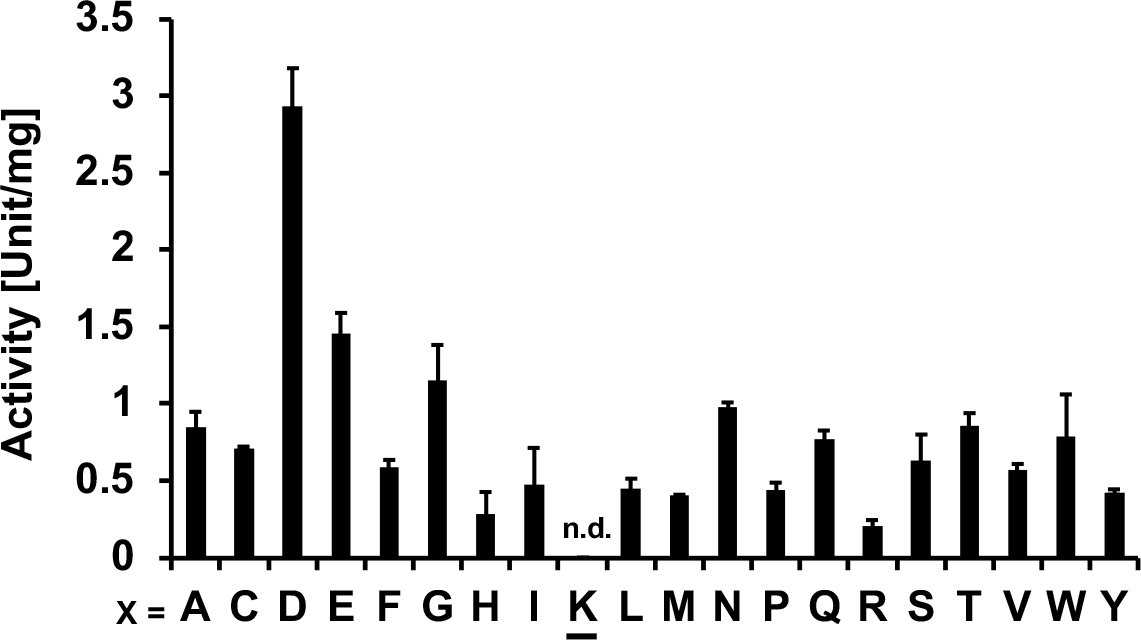
Enzymatic activity assay of K10X/Y12A MTGz mutants by a hydroxamate assay. The underlined K denotes the Y12A single mutant (n.d.: Not detected, below the detection limit). Data are presented as the mean ± standard deviation of three experiments (*n* = 3).

In terms of the protein expression, all mutants were found in the soluble fraction although each mutant showed different yield, implying the variation of chaperone functionality of mutated propeptides (**Fig. S5**). Notably, K10X/Y12A (X = D, E, G) mutants showed relatively lower protein yields but higher catalytic activity than other mutants. This trade-off relationship between the expression level of soluble proteins and their catalytic activity is likely due to cytotoxicity caused by the lack of activity regulation of MTG mutants in the *E. coli* cytoplasm. The yield of Y12A single mutant markedly decreased possibly due to the self-crosslinking and/or intermolecular crosslinking with endogenous proteins in the purification steps.

### Expression and enzymatic activity of K10R/Y12X mutants

We next focused on the position of Y12 with the combination of K10R mutation where K10 was replaced with Arg to minimize the effect of substitution by keeping the positive charge under the experimental conditions. A series of double mutants (K10R/Y12X MTGz, X = 19 natural amino acids) was expressed in *E. coli* and the enzymatic activity of purified enzymes was measured by a continuous GLDH-coupled assay (**Fig. S4 (ii)**).^41^ After the addition of MTG (0.15 μM) to the substrate solution containing cadaverine (primary amine substrate, 10 mM) and Z-QG (Gln-donor substrate, 20 mM), the catalytic activity was evaluated by the initial rates. K10R/Y12G mutant exhibited the highest activity (**Fig. 3**). Since Y12 constitutes a part of α-helical structure of the propeptide, the substitution with Gly decreased the stability of α-helix and may increase the accessibility of the substrate with the active site. The Y12D mutation also resulted in higher catalytic activity than the other mutants. Both of these mutants showed low protein yields in the expression (**Fig. S5**), and the trade-off relationship was shown as observed for K10X/Y12A mutants. Much lower yields of the K10R/Y12P mutant indicates that the substitution of Y12 with Pro, known to disrupt α-helix structure, would significantly reduce the intramolecular chaperone functionality of the propeptide domain.

**Fig. 3.**
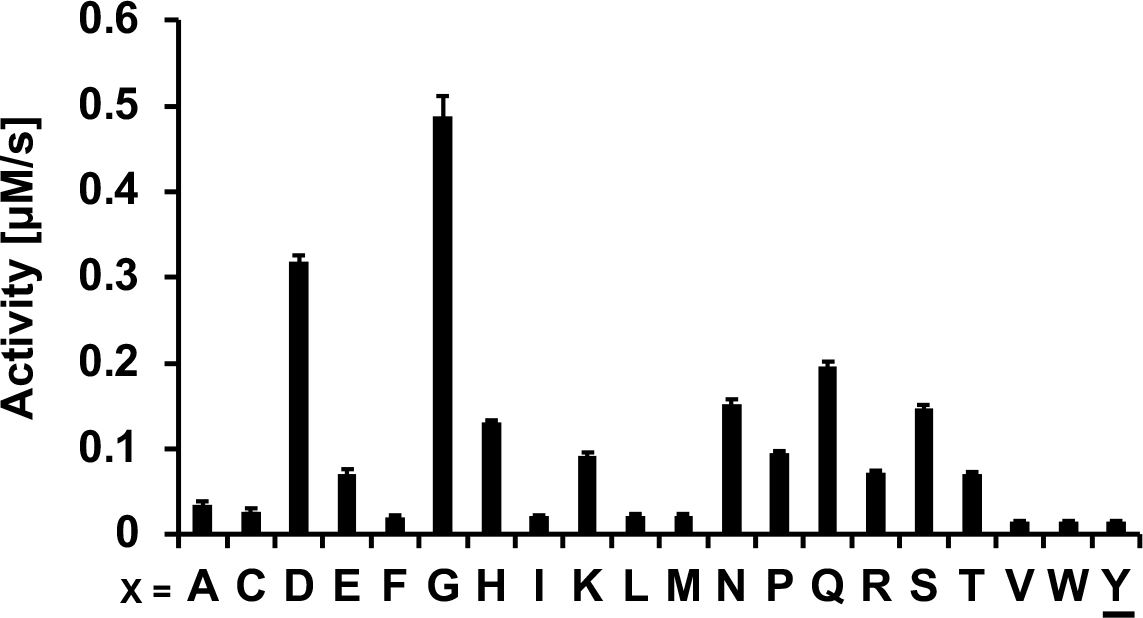
Enzymatic activity of K10R/Y12X mutants assessed by a continuous GLDH-coupled assay for MTG activity. The underlined Y denotes the K10R single mutant Data are presented as the mean ± standard deviation of three experiments (*n* = 3).

### Characterization of K10X_1_/Y12X_2_ double mutants

Encouraged by the results with K10X/Y12A and K10R/Y12X MTGz mutants, we investigated combined mutation at K10/Y12 with R/D for K10 and A/G for Y12, respectively. We first compared the susceptibility of two mutants, K10R (single point mutation) and K10R/Y12A mutants to thrombin digestion. SDS-PAGE analysis of the purified mutants before and after the thrombin treatment suggested that the propeptide was cleaved off from the zymogen scaffold by the substitution at Y12. By contrast, K10R single mutation was found to be resistant to the thrombin treatment (**Fig. 4A**). As expected, K10R mutant was unable to be fully activated by thrombin digestion whereas the specific activity of K10R/Y12A mutant was fully recovered to the level of ΔproMTG after thrombin digestion (**Fig. 4B**). These results suggest that the importance of Y12 position to control the susceptibility to the linker containing thrombin digestive site, which would also reflect the conformational flexibility around the propetide.

**Fig. 4.**
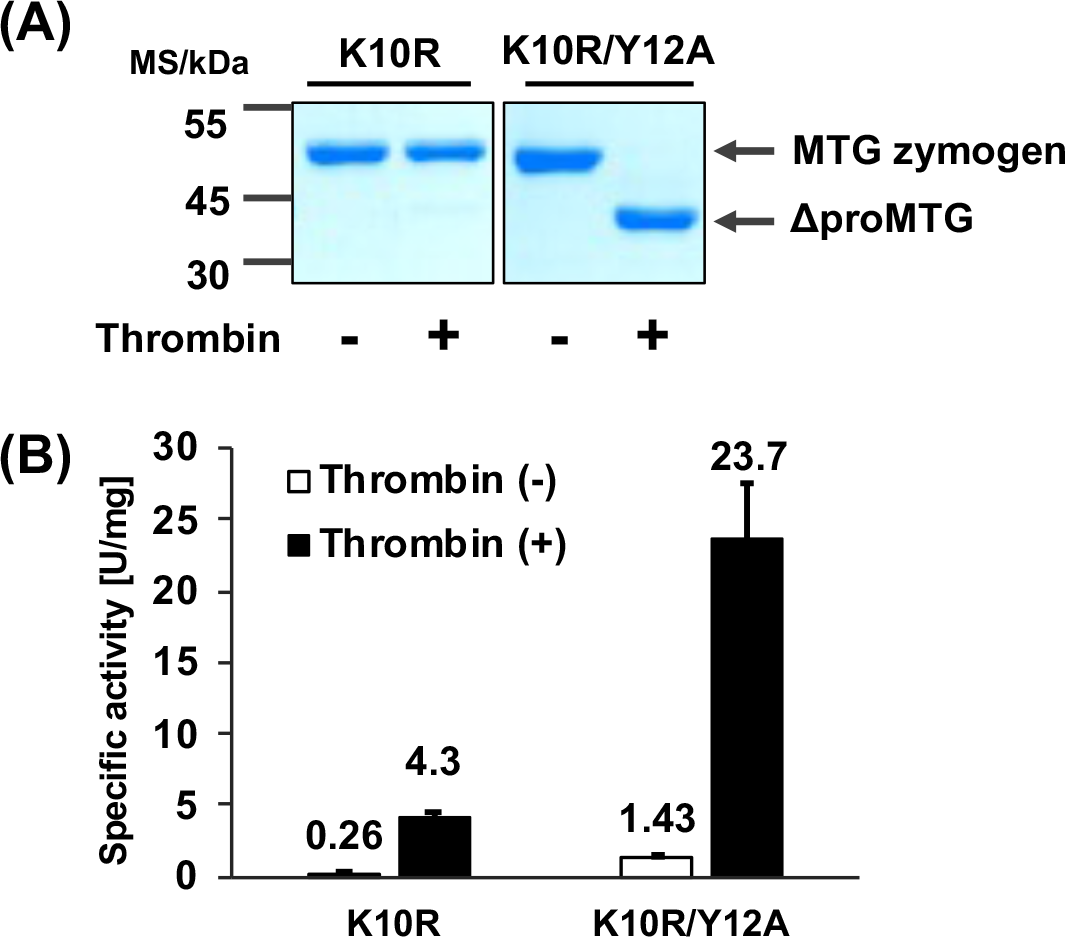
(A) SDS-PAGE analysis of purified K10R and K10R/Y12A MTGz mutants. (B) Specific activity of the mutants assessed by hydroxamate assay before and after thrombin digestion. Data are presented as the mean ± standard deviation of three experiments (*n* = 3).

Next we conducted the detailed kinetic analyses of four different types of EzMTG. From the inspection of the relationship between the initial rates and substrate concentration at diluted conditions, specificity constants (*k*_cat_/*K*_m_) were determined from the initial slope and were shown in **Fig. 5A**. We can see from the results that Y12G mutation had more impact on the increase in crosslinking activity than Y12A mutation for both of K10R and K10D double mutants. The difference was markedly enhanced by the integration of mutation at K10 position, and interestingly K10D/Y12G exhibited comparable specificity constant with that of ΔproMTG.

**Fig. 5.**
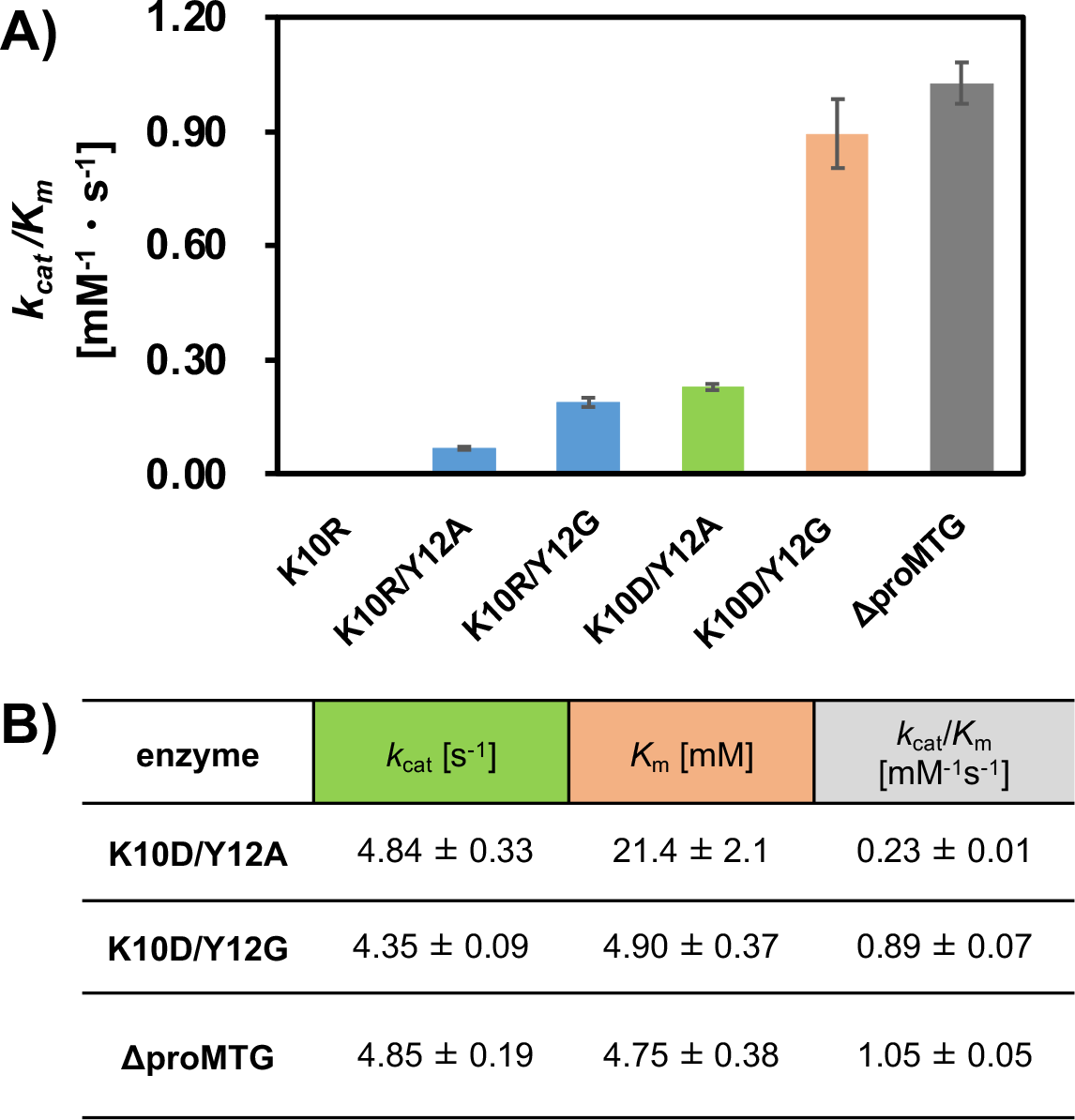
(A) Specificity constants (*k*_cat_/*K*_m_) of EzMTGs and ΔproMTG. (B) Kinetic parameters determined by using 2.5-30 mM Z-QG, 10 mM cadaverine and 0.15 μM enzyme in 0.2 M MOPS-EDTA (pH 7.2). Data are presented as the mean ± standard deviation of three experiments (*n* = 3).

For further analysis of the substrate recognition by MTGz mutants, two mutants were selected and kinetic parameters for Gln-donor substrate (Z-QG) were determined (**Fig. 5B**). It is revealed that K10D/Y12G mutant gained ca. 85% of the specificity constant of that of ΔproMTG and showed the comparable *K*_m_ value even in the presence of propetide in the protein scaffold. By contrast, K10D/Y12A mutant showed ca. 80% decrease in the specificity constant, mainly owing to ca. 5-fold increase in the *K*_m_ value. The results suggest that K10D/Y12A mutant is more susceptible to steric hindrance by propeptide upon the substrate binding in comparison to K10D/Y12G mutant.

The stability of enzymes is a crucial factor from the practical standpoint. Indeed, several critical point mutations to increase the thermal stability of mature MTG were reported.^42,43^ In terms of EzMTG, we found that the significant decrease in the catalytic activity of K10D/Y12G mutant upon the thermal treatment (**Fig. S6**). By contrast, K10R and K10R/Y12A mutants showed comparable thermal stability to ΔproMTG. These results imply the potential of MTGz mutants showing low activity as a target scaffold of EzMTG for protein engineering.

### Binding affinity of mutated propeptide to the enzyme active site

To gain further insight into the effects of mutated propetide on the substrate recognition by EzMTG, we designed fusion proteins composed of EGFP and mutated propetide, EGFP-Pro(K10X_1_/Y12X_2_), and they were subjected to bio-layer interferometry (BLI) analysis (**Fig. S7**).^44,45^ To eliminate the non-specific binding to the Ni-NTA-modified surface for BLI, ΔproMTG lacking the C-terminal His-tag (**Fig. S1**) was prepared and used as the analyte (**Fig. 6A**).

**Fig. 6.**
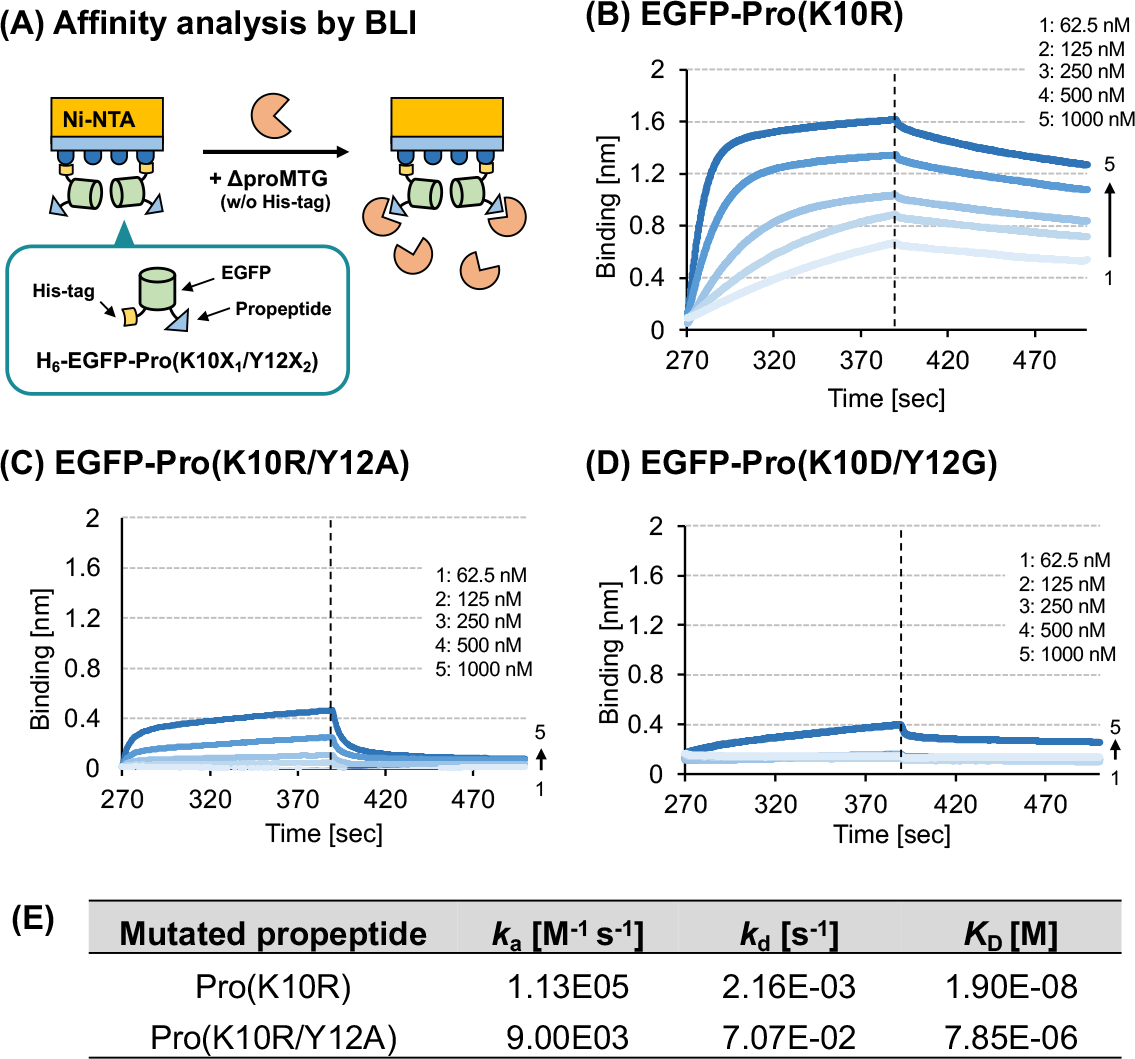
Evaluation of binding affinity of mutated propetide to the active site of MTG by bio-layer interferometry (BLI) analysis. (A) Schematic illustration of BLI analysis. (B)-(D) Association and dissociation profiles of ΔproMTG without His-tag to EGFP-Pro(K10R) or EGFP-Pro(K10R/Y12A) immobilized on the detection surface via Ni-NTA interaction. (E) Rate constants for association (*k*_a_) and dissociation (*k*_d_) and dissociation constant (*K*_D_) determined from the profiles of (B) and (C).

Based on the foregoing observations, K10R, K10R/Y12A and K10D/Y12G MTGz mutants showing different catalytic performance (**Fig. 5A**) were selected for the analysis. As expected, EGFP-Pro(K10R) showed the high affinity (*K*_d_ = 19.0 nM) with ΔproMTG (**Figs. 6B** and **6E**), by which little catalytic activity of K10R mutant can be explained. Interestingly, the additional mutation of Y12 to Ala resulted in ca. 410-fold reduction in the affinity (*K*_d_ = 7.85 μM) (**Figs. 6C** and **6E**), yet very low catalytic activity was observed for the K10R/Y12A mutant. These results indicate that the binding affinity of mutated propetide to the enzyme active site overrides the binding affinity of small molecular substrates, leading to the marked reduction of crosslinking activity. On the other hand, quantitative analysis with EGFP-Pro(K10D/Y12G) was impossible (**Fig. 6D**), indicating the disruption of tertiary structure of the propeptide, accounting the comparable catalytic activity to ΔproMTG and the low protein yield of K10D/Y12G MTGz mutant. These results demonstrated again that we should take both chaperone functionality and active site-binding affinity of mutated propeptide into account for the design of EzMTG.

Blood coagulation Factor XIII and TG2 are representative example of mammalian TGase in which the catalytic activity is controlled by endogenous Ca^2+^ ions. Mammalian TGase consists of several domains that undergo large conformational change upon the activation.^46,47,48^ Alternatively, our results toward EzMTG demonstrated that introduction of key mutations in propeptide domain could relieve the strong binding of the propeptide with the enzyme active site, enhancing the entry of substrates, thereby generating crosslinking activity with MTG zymogen.

### Application of EzMTG for site-specific protein labeling

Finally, we explored the potential of EzMTG in protein labeling applications (**Fig. S4 (iii)**). **Fig. 7** shows the SDS-PAGE analysis of cross-linking reaction between TNF-α fused with the N-terminal MRHKGS sequence (Ktag-TNF-α) and FITC-β-Ala-QG. The fluorescent images showed that Ktag-TNF-α was successfully labeled with the fluorescent probe by ΔproMTG and all the EzMTGs except for K10R mutant. Time-dependent increase in the fluorescent signal revealed that the trend of crosslinking reaction, which was consistent with the order of catalytic activity found in **Fig. 5**. Notably, K10R/Y12A MTGz mutant showed > 50% yield after prolonged incubation for 1 h, though this mutant exhibited ca. 1/10-fold crosslinking activity for small molecular substrates (**Fig. 5A**). The results imply that the substrate recognition of EzMTG depends on the type of substrates, namely the difference in the affinity of substrate to the active site of MTG. This also suggests that the substrate preference can be further tuned toward a specific substrate by designing a high-throughput screening system with EzMTG because soluble expression of active mutants in cytosol is possible, leading to further manipulation of MTG and possibly to extend our concept to other precursor enzymes.

**Fig. 7.**
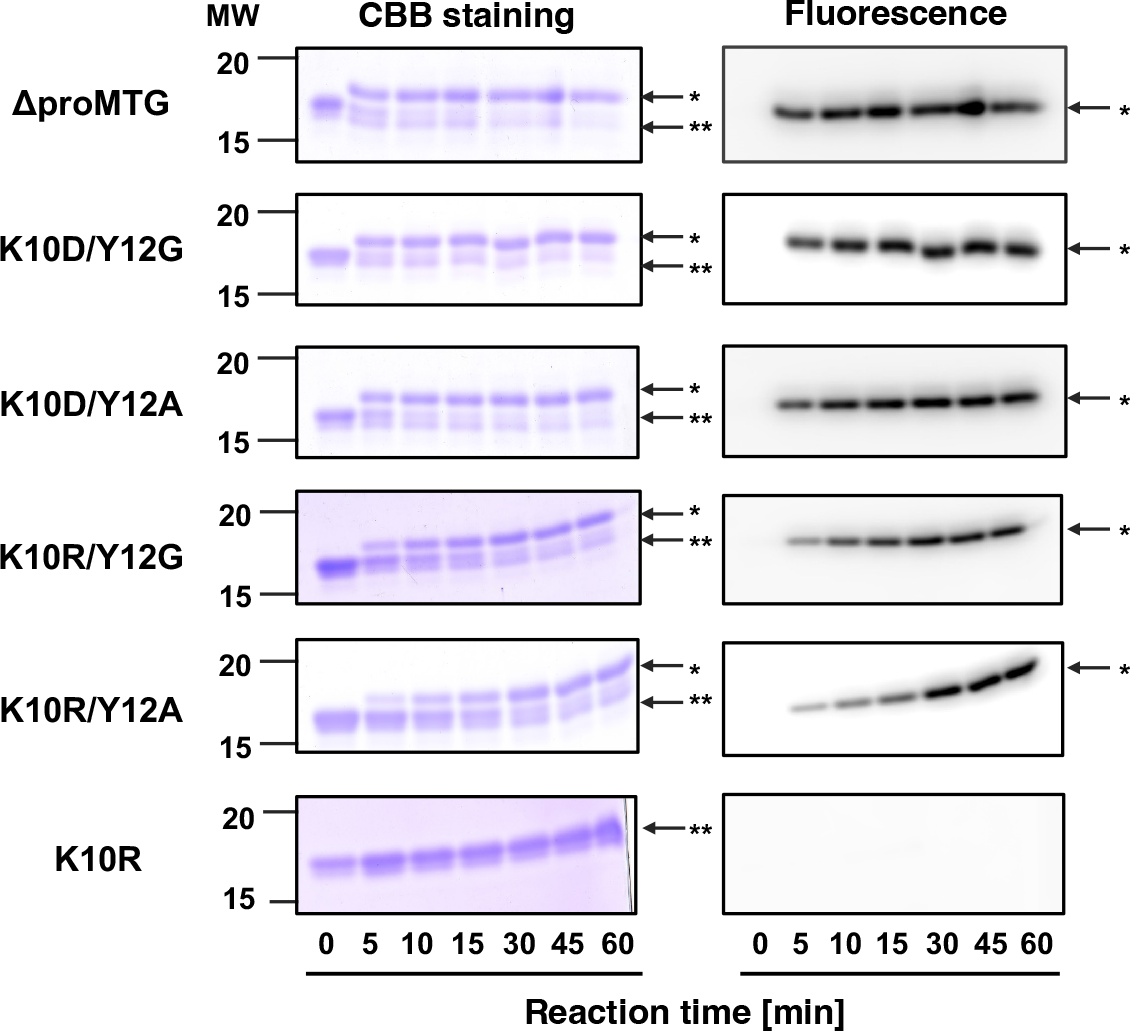
SDS-PAGE analysis of ΔproMTG- or EzMTG-mediated labeling of Ktag-TNFα with FITC-β-Ala-QG. The results of Coomassie Brilliant Blue staining (left) and the fluorescence (right) of the gel. *FITC-labeled Ktag-TNFα, **Non-labeled Ktag-TNFα.

## Conclusion

Our concept of EzMTG proposes one straightforward way to promote the activation of MTG zymogen without limited proteolysis by either inducing conformational fluctuation in propeptide domain or keeping the chaperone functionality. Our proof-of-concept study will lead to generate comparable catalytic performance with altered substrate specificity to mature MTG as exemplified by K10R/Y12A MTGz mutant that exhibited an improved crosslinking activity for peptidyl substrates. Unique substrate preference by modulating the intramolecular inhibitory domain of an enzyme will provide insight into the design of a new biocatalyst from its precursor. Since EzMTG can be expressed as soluble form in the *E. coli* cytosol, our results will provide an opportunity for the high-throughput screening of active MTG zymogen mutants for *in vivo* bioconjugation applications.

## Materials and methods

### 1. Materials

The PrimeSTAR® MAX DNA Polymerase and In-Fusion® HD Cloning Kit were purchased from Takara Bio Inc. (Shiga, Japan). LB broth (Miller) was purchased from Merck Millipore Corp (Billerica, MA, USA). Tryptone, dried yeast extract, tris(hydroxymethyl)aminomethane (Tris) and the Rapid Stain CBB Kit were purchased from NACALAI TESQUE, INC (Kyoto, Japan). Hydrochloric acid, sodium chloride, potassium chloride, disodium hydrogen phosphate 12-water, dipotassium hydrogen phosphate and potassium dihydrogen phosphate were purchased from FUJIFILM Wako Pure Chemical Corporation (Osaka, Japan). The 5∼20% gradient gel for SDS-PAGE was purchased from ATTO CORPORATION (Tokyo, Japan). N-Ethylmaleimide was purchased from KISHIDA CHEMICAL Co., Ltd. (Osaka, Japan). Ultrapure water supplied by a Milli-Q® Reference Water Purification System (Merck Millipore Corp.) was used for buffer preparation. A matured MTG used as a control was purchased from Zedira (Darmstadt, Germany). MTG was recombinantly prepared by Escherichia coli BL21 star (DE3) as previously reported^49^.

### 2. Construction of MTG zymogen mutants

The pET22b+ vector carrying the gene encoding K10R/Y12A MTG^49^ was used as the template DNA for vector construction of K10X/Y12A and K10R/Y12X mutants (X = 20 natural amino acids). To construct the vector containing the MTG mutants, some mutations were introduced by inverse PCR and self-ligation of the PCR products, introducing a DNA sequence of thrombin cleavage sequence (LVPRGS) between the propeptide and mature MTG. The amino acid sequences and schematic illustrating of all the MTG mutants used in this study were summarized in **Table S1**.

### 3. Expression and purification of recombinant proteins

#### 3.1. K10X/Y12A and K10R/Y12X MTG mutants

The expression of the MTG mutants were conducted in an *Escherichia coli* BL21Star(DE3) strain. The plasmid vectors of MTG mutants were transformed into the cells by heat shock and cultured on a lysogeny broth plate containing 100 μg/mL of sodium ampicillin. A single colony was inoculated into 3.5 mL of lysogeny broth containing the same amount of sodium ampicillin and shaken at 220 rpm and 37°C for 4 h. The precultured medium of Pro (K10X/Y12A) MTG were placed into 25 mL of TB Medium containing 100 μg/mL ampicillin and cultured with shaking at 220 rpm at 37°C until the OD_600_ reached 0.5. The expression of K10X/Y12A MTG mutants were induced by the addition of 1 mM isopropyl-β-D-thiogalactoside (IPTG) and followed by shaking at 18 °C. K10R/Y12X MTG mutatns were cultured with Auto Induction Medium without inducer such IPTG. The cells were harvested by centrifugation at 5000 × g for 20 min and supernatant was removed. The pellet was resuspended in PBS (pH 7.4) and frozen at −80°C until purification.

For the purification of MTG mutants, the pellet was thawed in running water. The cell suspension was sonicated on ice for 3 min. The cell debris was removed by centrifugation at 16, 000 × *g* for 10 min at 4 °C. The supernatants were initially applied to a HisGraviTrap column (GE Healthcare Life Sciences, Chicago, IL, USA), which was pre-equilibrated with HisTrap binding buffer (20 mM Tris-HCl, 500 mM NaCl, 35 mM imidazole, pH 7.4). The columns were washed with the same buffer until all of the unbound proteinaceous substances were eluted, and then the peptide tag fused proteins were eluted with a HisTrap elution buffer (20 mM Tris-HCl, 500 mM NaCl, 500 mM imidazole, pH 7.4). The fractions containing MTG variants were concentrated, and the buffer was exchanged to binding buffer (1×PBS with 0.1 mM Dithiothreitol, pH 7.4) after membrane filtration using a 10 kDa MWCO (Amicon Ultra-15 Centrifugal Filter Units, Billerica, MA, USA). The purified MTG mutants were concentrated using the ultrafiltration membrane that was suitable for each protein. Final protein concentration of MTG mutants was quantified by YabGelImage software (https://sites.google.com/site/yabel/home) using bovine serum albumin (BSA) as a standard protein.

#### 3.2. K10D/Y12G MTG mutant

The expression of Pro (K10D/Y12G) MTG was conducted on an *Escherichia coli* BL21 Star (DE3) strain. The plasmid vectors of the MTG mutant was transformed into the cells by a heat shock method and cultured on a lysogeny broth plate containing 100 μg/mL of ampicillin sodium. A single colony was inoculated to 3.5 mL of lysogeny broth containing the same amount of ampicillin sodium and shaken at 220 rpm and 37 °C 6 hours. The precultured medium was placed into 1 L of Terrific broth containing the same antibiotics and cultured with shaking at 220 rpm at 37 °C until the optical density at 600 nm reached around 0.5. Expression of each protein was induced by addition of isopropyl β-d-1-thiogalactopyranoside (IPTG) (final concentration of 0.1 mM), and the cells were cultured overnight after lowering the temperature to 17 °C. The cells were harvested by centrifugation at 5, 000×g for 20 min and supernatant was removed. Pellet was resuspended by PBS (pH 7.4) and frozen at −80 °C until purification.

For the protein purification, the pellet was thawed with running water. The cell suspension was sonicated on ice for 12.5 min. The cell debris was removed by centrifugation at 15, 000 × g for 10 min at 4 °C. The supernatants were initially applied to a HisTrap FF crude 5 mL (GE Healthcare UK Ltd.), which were pre-equilibrated with the HisTrap binding buffer (20 mM Tris-HCl, 500 mM NaCl, 35 mM imidazole, pH 7.4). The columns were washed with the same buffer until all of the unbound proteinaceous substances eluted out, and MTG mutant was eluted by a gradient of HisTrap elution buffer (20 mM Tris-HCl, 500 mM NaCl, 500 mM imidazole, pH 7.4) up to 100% with 13 column volumes. The fractions containing MTG mutant was concentrated, and the buffer was exchanged to HiTrap binding buffer (10 mM Tris-HCl, pH 8.0) after membrane filtration of 10 kDa MWCO (Amicon Ultra-15 Centrifugal Filter Units, Billerica, MA, USA). The desalted solution containing protein of interest was applied to a HiTrap Q HP 5 mL column (GE Healthcare UK Ltd.) pre-equilibrated with the HiTrap binding buffer. The column was washed with five column volumes of the same buffer, and peptide tag fused proteins were eluted by a gradient of elution buffer (10 mM Tris-HCl, 500 mM NaCl, pH 8.0) up to 100% with 13 column volumes. The fractions containing peptide tag fused proteins were concentrated to a volume of approximately 5 mL using the ultrafiltration membrane, and MTG mutant was purified with size exclusion chromatography using ProteoSEC-D 16/60 6-600 HR (Protein Ark) with PBS (pH 7.4) including 0.1 mM dithiothreitol as a running buffer. The purified protein was concentrated using the ultrafiltration membrane which was suitable for MTG mutant. To determine protein concentrations, the absorbance at 280 nm was measured using an ND-1000 Nanodrop (NanoDrop) and protein concentration was calculated using the extinction coefficient of each mutant predicted by ExPAsy ProtParam (https://web.expasy.org/protparam/).

#### 3.3 EGFP-Propeptide fusion proteins

For the purification of EGFP-propeptide mutants, the pellet was thawed in running water. The cell suspension was sonicated on ice for 3 min. The cell debris was removed by centrifugation at 15, 000 × g for 10 min at 4°C. The supernatants were initially applied to a HisGraviTrap column (GE Healthcare Life Sciences, Chicago, IL, USA), which was pre-equilibrated with HisTrap binding buffer (20 mM Tris-HCl, 500 mM NaCl, 35 mM imidazole, pH 7.4). The columns were washed with the same buffer until all of the unbound proteinaceous substances were eluted, and then the peptide tag fused proteins were eluted with a HisTrap elution buffer (20 mM Tris-HCl, 500 mM NaCl, 500 mM imidazole, pH 7.4). The fractions containing EGFP-propeptide mutant was concentrated, and the buffer was exchanged to HiTrap binding buffer (10 mM Tris-HCl, pH 8.0) after membrane filtration of 10 kDa MWCO (Amicon Ultra-15 Centrifugal Filter Units, Billerica, MA, USA). The desalted solution containing protein of interest was applied to a HiTrap Q HP 5 mL column (GE Healthcare UK Ltd.) preequilibrated with the HiTrap binding buffer. The column was washed with five column volumes of the same buffer, and peptide tag fused proteins were eluted by a gradient of elution buffer (10 mM Tris-HCl, 500 mM NaCl, pH 8.0) up to 100% with 13 column volumes. The fractions containing EGFP-propeptide mutant was concentrated, and the buffer was exchanged to PBS (pH 7.4) after membrane filtration of 10 kDa MWCO. To determine protein concentrations, the absorbance at 280 nm was measured using an ND-1000 Nanodrop (NanoDrop).

#### 3.4 K-tagged TNF-α

The synthetic gene encoding matured human TNF-α with an N-terminal K-tag sequence (MRHKGS) synthesized by and purchased from IDT Technologies was amplified by PCR and cloned into the pCold 4 vector using In-Fusion Cloning to construct the expression vector for K-tagged TNF-α (Ktag-TNFα). The vector was transformed into competent cells of E. coli BL21star (DE3) using the heat shock method (42°C for 30 seconds) and cultured on an LB agar plate containing 100 μg/mL of ampicillin sodium. After incubating overnight at 37°C, a single colony was picked and inoculated into 5 mL of LB medium containing the same amount of antibiotics and cultured overnight at 37°C with shaking at 180 rpm. The precultured cell suspension was then transferred into 1 L of LB medium containing 100 μg/mL of ampicillin sodium and cultured at 37°C with shaking at 180 rpm until the OD600 reached 0.5. The cultured cell suspension was then cooled on ice for approximately 10 minutes to reduce the temperature to below 17°C. Then, IPTG was added at a final concentration of 0.1 mM and the culture was continued for 12 hours at 17°C. The cells were collected by centrifugation at 5,000 G for 20 minutes. The cells were washed with PBS(-) twice and frozen at -80 °C until purification. The cells were then thawed and lysed by sonication for 3 minutes, three times, with a 3-minute interval for each cycle. The cell lysate was centrifuged at 15,000 G for 30 minutes to separate the insoluble debris. The supernatant was filtered through a 0.45 μm membrane filter and then applied to a 5 mL HisTrap FF crude column, which was pre-equilibrated with His-Binding buffer (25 mM Tris HCl pH 7.4, 500 mM NaCl, 25 mM imidazole). The column was washed with His-Binding buffer until the absorbance of the column eluate at 280nm stabilized, and then the Ktag-TNFα was eluted from the column using a gradient of the His-Elution buffer (25 mM Tris HCl pH 7.4, 500 mM NaCl, 500 mM imidazole) from 0% to 100% over a volume of 25 mL. The fractions containing proteins were analyzed by SDS-PAGE to confirm the presence of the Ktag-TNFα. The fractions containing Ktag-TNFα were collected and subjected to buffer exchange into 10 mM Tris HCl pH 7.4 using a HiPrep 26/10 Desalting column. The buffer-exchanged solution was then applied to a HiTrap SP HP column pre-equilibrated with 10 mM Tris HCl pH 7.4. The column was washed with 25 mL of the same buffer, and then Ktag-TNFα was eluted using a gradient of the same buffer containing 500 mM NaCl from 0% to 100%. The fractions containing the target protein were collected and concentrated to about 5 mL using an ultrafiltration membrane (10 kDa MWCO). The concentrated solution was applied to a size exclusion column (HiLoad 16/600 Superdex 200 pg) that had been pre-equilibrated with 10 mM Tris-HCl (pH 8.0) and further purified by size exclusion chromatography. The fractions were analyzed with SDS-PAGE. The fractions containing Ktag-TNFα were collected and concentrated to approximately 3 mL using an ultrafiltration membrane (10 kDa MWCO) and stored at -80°C in aliquots. The concentration of the purified Ktag-TNFα was determined by measuring the UV absorbance at 280 nm using NanoDrop.

### 4. Activity measurement of the MTG mutants

#### 4.1 Hydroxamate assay

K10X/Y12A MTG mutants were assessed by quantifying the amount of hydroxamate resulted from MTG reaction using complex formation of hydroxamate and Fe(III) ions^50^. The substrate solution containing 30 mM Z-QG, 100 mM hydroxylamine hydrochloride, 10 mM glutathione in 0.2 M Tris-Acetate (pH 6.0) was added to MTG variants. After 10 min of incubation at 37 °C, the stop solution (200 mM trichloroacetic acid in 50 mM HCl) containing 50 mM of Fe(III) chloride hexahydrate was added to the reaction mixture and the absorbance at 525 nm was measured using Synergy HTX Multi-Mode Reader (BioTek, Winooski, USA). A standard curve was made by using L-glutamic acid γ-monohydroxamate solution with known concentration. One unit of MTG was defined as the amount of MTG catalyzes formation of 1 μmol of hydroxamic acid in one minute.

### 4.2 GLDH-coupled assay

To determine the activity of K10R/Y12X MTG mutants, a continuous GLDH-coupled assay for MTG activity was applied^41^. For glutamine substrate evaluation, the assay was performed in a 96-well half plate in the presence of 10 mM α-ketoglutarate, 2 U of glutamine dehydrogenase (GLDH), 500 μM NADH, 10 mM cadaverine as amine donor substituting for a Lys peptide, 20 mM Z-Gln-Gly as acyl-acceptor substituting for a Gln peptide, 1 mM EDTA and 200 mM MOPS buffer (pH 7.2) in a final volume of 90 μL. The reaction solution was preincubated for 10 min at 37 °C. Reaction were started by the addition of 1.5 μM Pro (K10R/Y12X) MTG (in a volume of 10 μL) to give a final volume of 200 μL, and the oxidation of NADH was continuously recorded at 340 nm for 60 min using a Microplate reader (SpectraMax i3x, Molecular Device), temperature-controlled at 37 °C, with short shaking intervals before each measurement. After a short lag phase during which sufficient ammonia is produced to saturate GLDH, linear slopes of absorbance versus time were measured, then the active of MTG were determined by initial rates of NADH.

For the kinetics analysis EzMTG and active mature MTG (ΔproMTG)^48^, initial reaction rates were measured by varying the concentration of Z-QG at the constant concentration of cadaverine. Initial rates data are fitted to Michaelis-Menten kinetics, then kinetic parameters were determined by KaleidaGraph™ software.

#### 4.3 Thermal stability measurements

Thermal stability of selected EzMTG mutants were evaluated by incubating the aqueous solution of the enzyme in PBS buffer (pH 7.4) at 60 °C for 1 h.

### 5. Binding affinity of mutated propeptide to the MTG active site

Binding kinetics for EGFP-Propeptide and MTG were measured using a biolayer interferometry system (BLItz) with Ni-NTA biosensors (ForteBio Inc.). Sensor tips were prehydrated for 10 min in Milli-Q and then equilibration in K buffer (150 mM NaCl, 0.1% BSA, 0.002% Tween20). 100 ug/mL of EGFP-propeptide was immobilized to Ni-NTA sensor tips for 2 min during the binding of proteins reached saturation. A concentration series of 62.5, 125, 250, 500, 1000 nM of ΔproMTG was allowed for 2 min followed by 2 min dissociation step. We tested 100 ug/mL of WT-EGFP as negative control and 1000 nM of ΔproMTG was used. Assays were performed according to the instrument manual. Data were exported from BILtz pro software and replotted. Values of binding as reflected by changing in optical thickness (nm) were used to calculate each kinetic constants (*ka, kd* and *K*_*D*_) using nonlinear curve fitting for entire concentration of ΔproMTG.

### 6. MTG-mediated labeling of K-tag TNF-α with fluorescent substrates

The reaction mixture comprised TNF-α-Ktag (10 μM) and FITC-β-Ala-QG (200 μM) in PBS buffer (pH 7.4)^51^. The protein labeling reaction was initiated by the addition of MTG (0.1 μM) at 37 °C. To follow the time course of MTG-mediated labeling of TNF-α-Ktag with FITC-β-Ala-QG, a small aliquot of 4 × SDS-PAGE sample buffer was added to the reaction solution to terminate the MTG reaction. The reaction was followed by the increase in the fluorescence of the protein bands in the fluorescent image of the SDS-PAGE gel. With respect to the fluorescein derivatives, 6 μL of the reaction samples were applied to the gel. Before staining the gel with Coomassie Brilliant Blue (CBB), the fluorescent image of the gels was obtained using a Molecular Imager FX Pro (Bio-Rad Laboratories, Inc.). An excitation wavelength of 488 nm with a 530 (±15) nm band pass filter for the fluorescein-derivatives was used.

## Acknowledgments

This work was supported by the Japan Society for the Promo-tion of Science (JSPS) KAKENHI Grant numbers JP19H00841 and JP23H00247 (to N. K.)

## Notes

### Competing Interest Statement

The authors have declared no competing interest.

