## Supporting information for "Engineered Active Zymogen of Microbial Transglutaminase"

**Table S1.** The amino acid sequences of all the proteins used in this study.

### 1-1. K10$X_1$/Y12$X_2$ MTGz mutants ($X_1$ and $X_2$ = natural amino acids)

MDNGAGEET$X_1$S$X_2$AETYRLTADDVANINALNESAPAASSGGGSLVPRGSGGGSAPDSDDRVTPPAEPLDRMPDPYRPS
YGRAETVVNNYIRKWQQVYSHRDGRKQQMTEEQREWLSYGCVGVTWVNSGQYPTNRLAFASFDEDRFKNELKNGRPRS
GETRAEFEGRVAKESFDEEKGFQRAREVASVMNRALENAHDESAYLDNLKKELANGNDALRNEDARSPFYSALRNTPS
FKERNGGNHDPSRMKAVIYSKHFWSGQDRSSSADKRKYGDPDAFRPAPGTGLVDMSRDRNIPRSPTSPGEGFVNFDYG
WFGAQTEADADKTVWTHGNHYHAPNGSLGAMHVYESKFRNWSEGYSDFDRGAYVITFIPKSWNTAPDKVKQGWPGSHH
HHHH

### 1-2. EGFP-Pro(K10$X_1$/Y12$X_2$) fusion ($X_1$ = R or D, $X_2$ = A or G)

MHHHHHHMVSKGEELFTGVVPILVELDGDVNGHKFSVSGEGEGDATYGKLTLKFICTTGKLPVPWPTLVTTLTYGVQC
FSRYPDHMKQHDFFKSAMPEGYVQERTIFFKDDGNYKTRAEVKFEGDTLVNRIELKGIDFKEDGNILGHKLEYNYNSH
NVYIMADKQKNGIKVNFKIRHNIEDGSVQLADHYQQNTPIGDGPVLLPDNHYLSTQSALSKDPNEKRDHMVLLEFVTA
AGITLGMDELGGGGSDNGAGEET$X_1$S$X_2$AETYRLTADDVANINALNESAPAASS

### 1-3. ΔproMTG (w/o $H_6$-tag)

SGGGGSDSDDRVTPPAEPLDRMPDPYRPSYGRAETVVNNYIRKWQQVYSHRDGRKQQMTEEQREWLSYGCVGVTWVNS
GQYPTNRLAFASFDEDRFKNELKNGRPRSGETRAEFEGRVAKESFDEEKGFQRAREVASVMNRALENAHDESAYLDNL
KKELANGNDALRNEDARSPFYSALRNTPSFKERNGGNHDPSRMKAVIYSKHFWSGQDRSSSADKRKYGDPDAFRPAPG
TGLVDMSRDRNIPRSPTSPGEGFVNFDYGWFGAQTEADADKTVWTHGNHYHAPNGSLGAMHVYESKFRNWSEGYSDFD
RGAYVITFIPKSWNTAPDKVKQGWP

### 1-4. Ktag-EGFP

MKHKGSMVSKGEELFTGVVPILVELDGDVNGHKFSVSGEGEGDATYGKLTLKFICTTGKLPVPWPTLVTTLTYGVQCF
SRYPDHMKQHDFFKSAMPEGYVQERTIFFKDDGNYKTRAEVKFEGDTLVNRIELKGIDFKEDGNILGHKLEYNYNSHN
VYIMADKQKNGIKVNFKIRHNIEDGSVQLADHYQQNTPIGDGPVLLPDNHYLSTQSALSKDPNEKRDHMVLLEFVTAA
GITLGMDELYHHHHHH

### 1-5. Ktag-TNF-α

MRHKGSGGHHHHHHGGGGSGGGGSVRSSSRTPSDKPVAHVVANPQAEGQLQWLNRRANALLANGVELRDNQLVVPSEG
LYLIYSQVLFKGQGCPSTHVLLTHTISRIAVSYQTKVNLLSAIKSPCQRETPEGAEAKPWYEPIYLGGVFQLEKGDRL
SAEINRPDYLDFAESGQVYFGIIAL

Gray: Propeptide domain; Red: Mutated amino acid; Black: mature MTG domain; Purple: Linker sequences; Orange: Hexahistidine tag; Blue: K-tag; Green: EGFP domain; Brown :TNF-α.

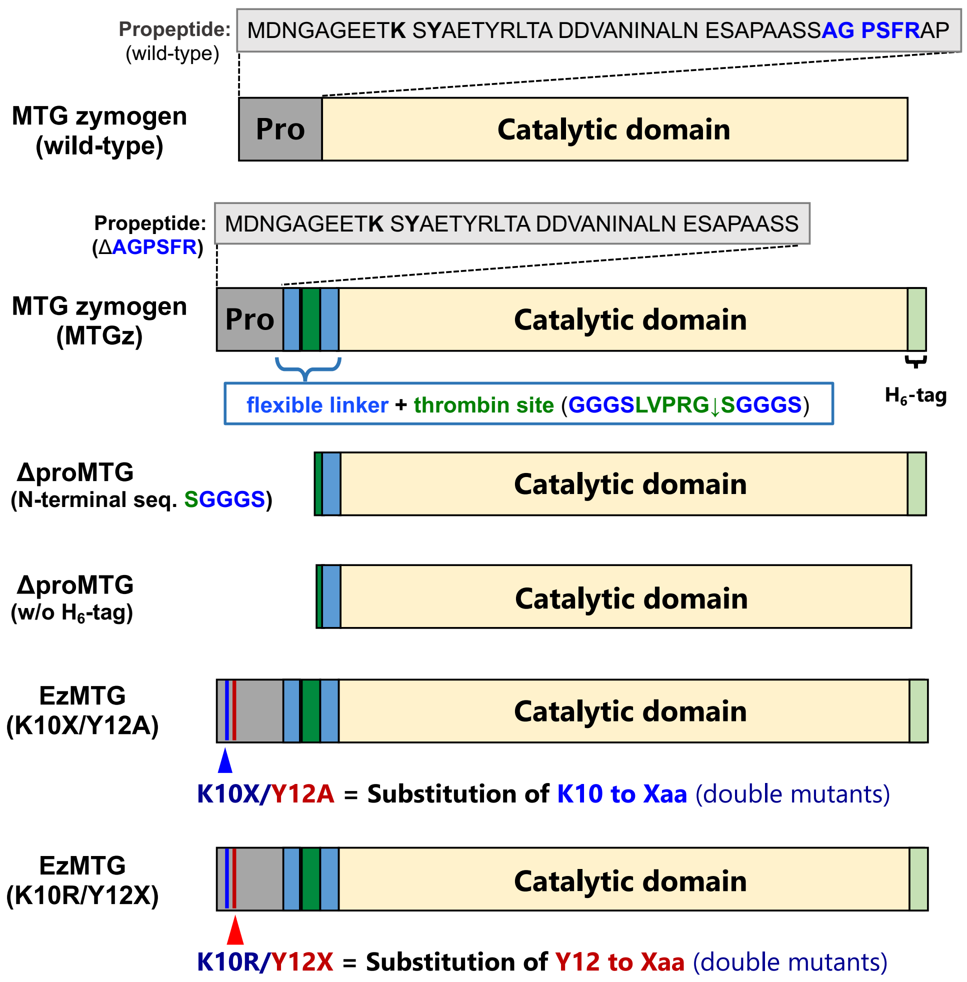


**Fig. S1** Designed MTG mutants with abbreviations prepared in this study.

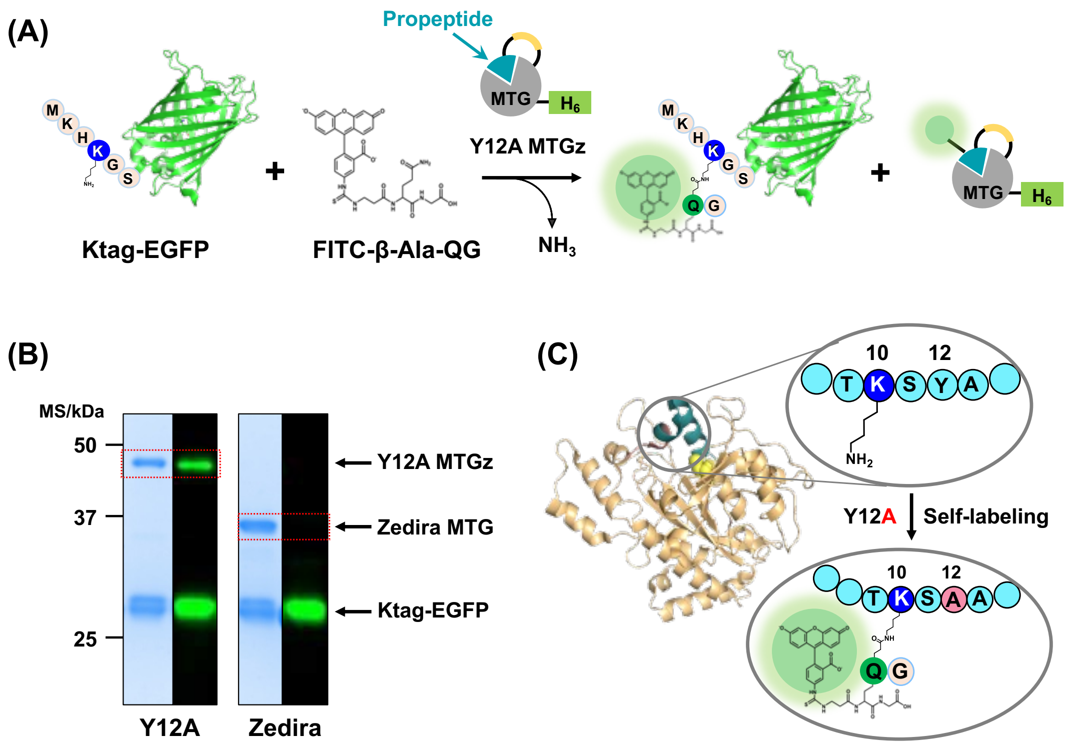


**Fig. S2** Verification of self-labeling of Y12A MTG zymogen. (A) Schematic illustration of the site-specific labeling of Ktag-EGFP and the self-labeling of Y12A MTGz with FITC-β-Ala-QG (B) SDS-PAGE and fluorescence analysis of the self-labeling of Y12A MTG and a commercial MTG (Zedira) (C) Schematic illustration of self-labeling of K10 in the propeptide.

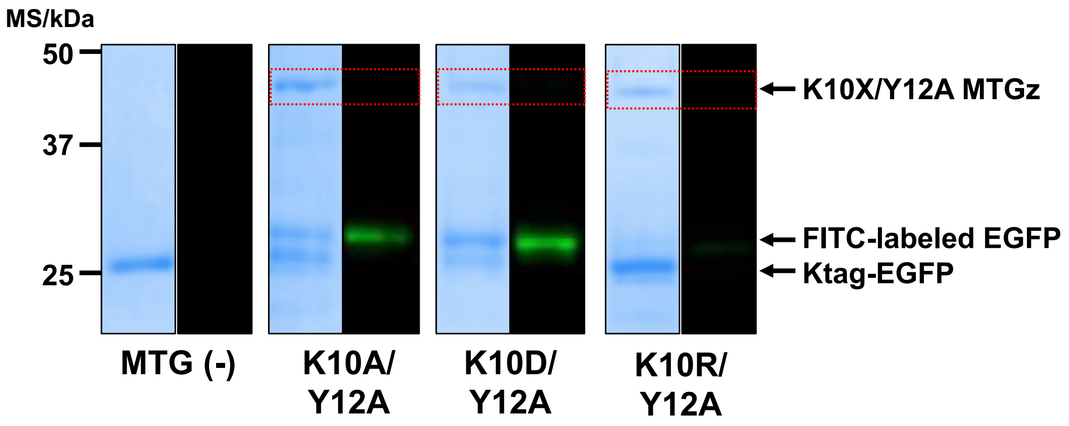


**Fig. S3** Elimination of self-labeling by site-directed mutagenesis of K10 in the propeptide.

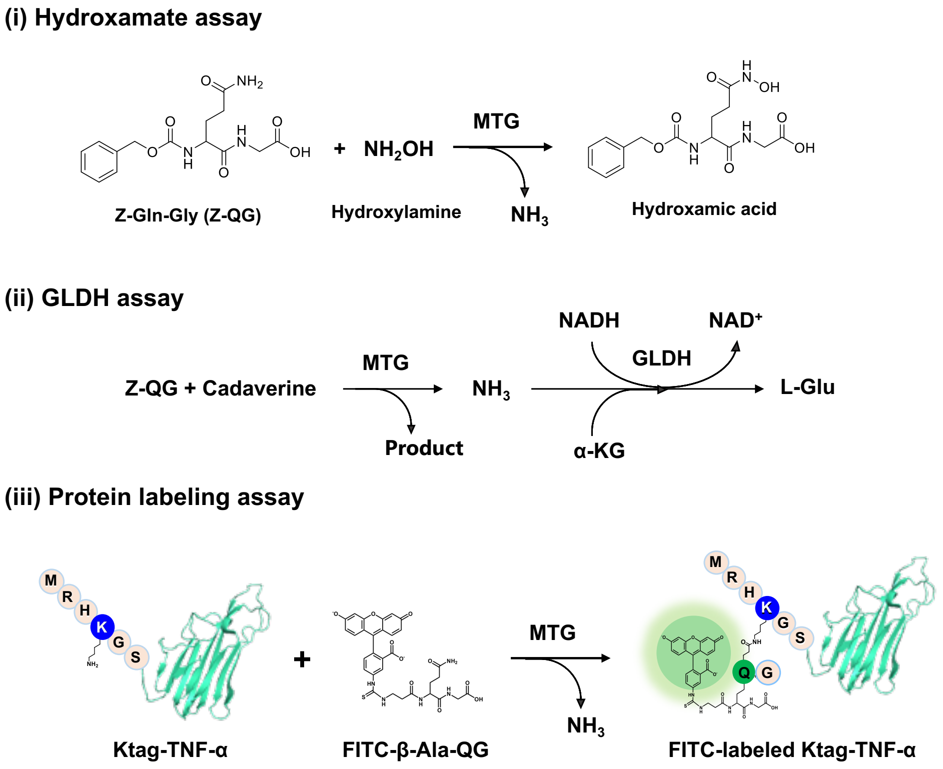


**Fig. S4** Scheme of enzymatic activity assays

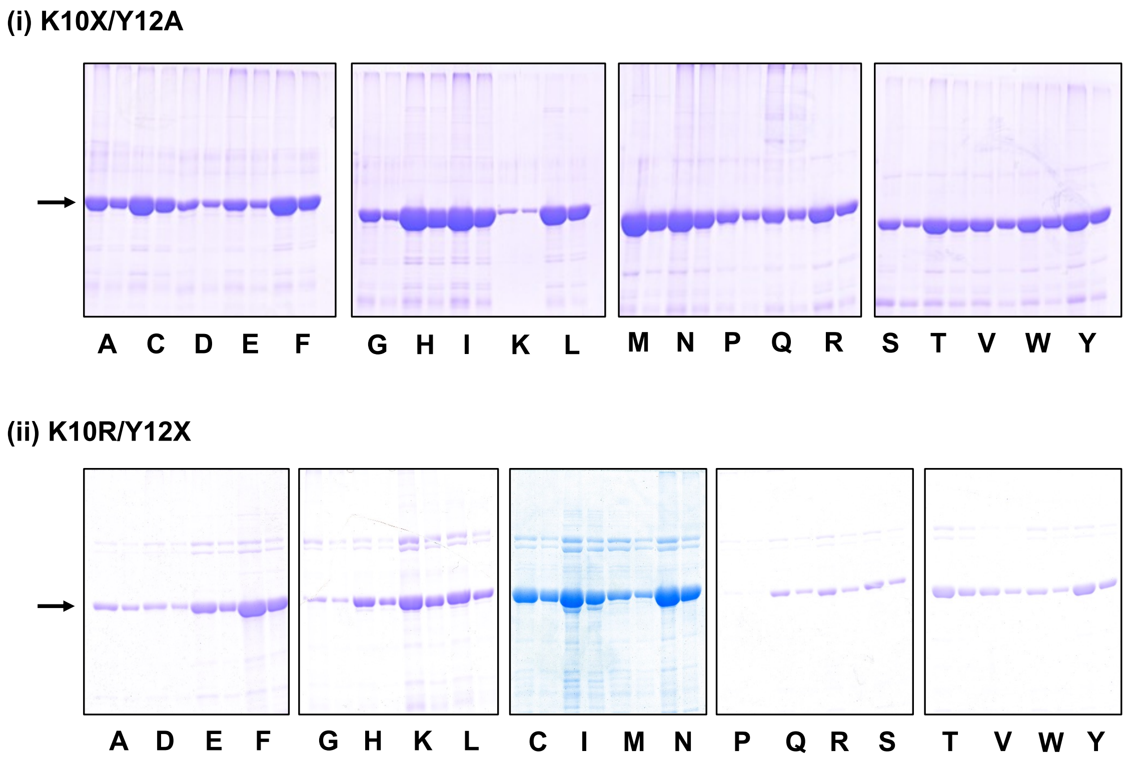


**Fig. S5** Results of the expression of K10$X_1$/Y12$X_2$ MTGz mutants in *E. coli*. SDS-PAGE images showed EzMTG samples at different dilution rates to quantify the expression level.

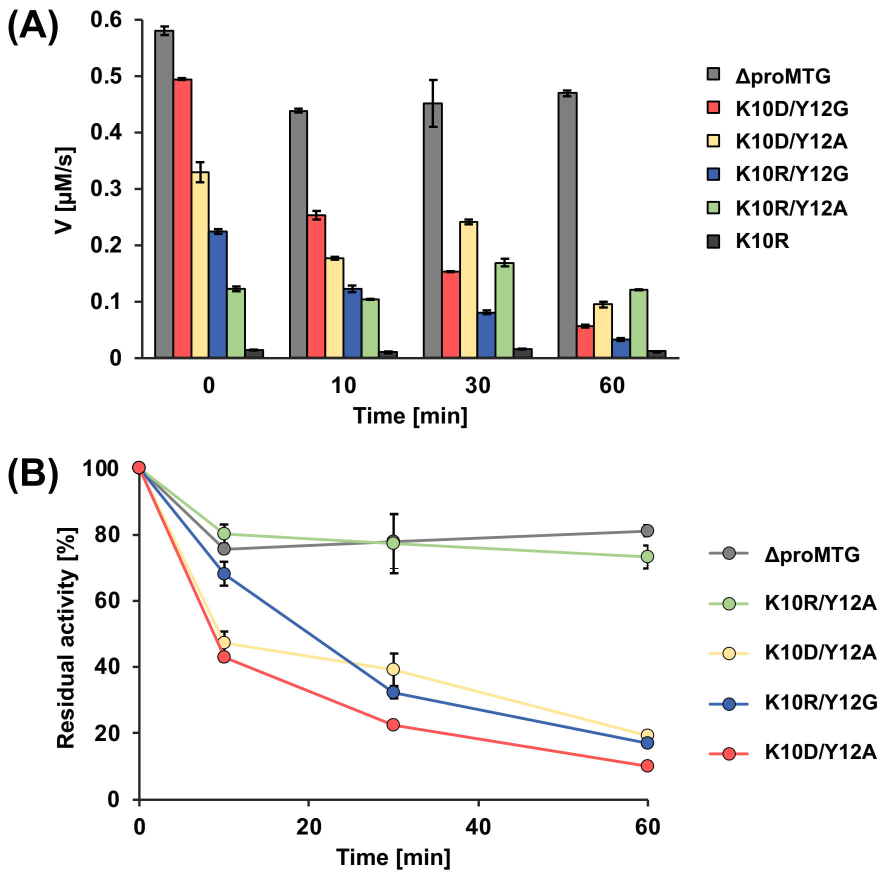


**Fig. S6** (A) Comparison of enzymatic activities upon thermal treatments at 60 °C for 1 h. (B) the time course of relative activity change of EzMTG mutants except for K10R mutant.

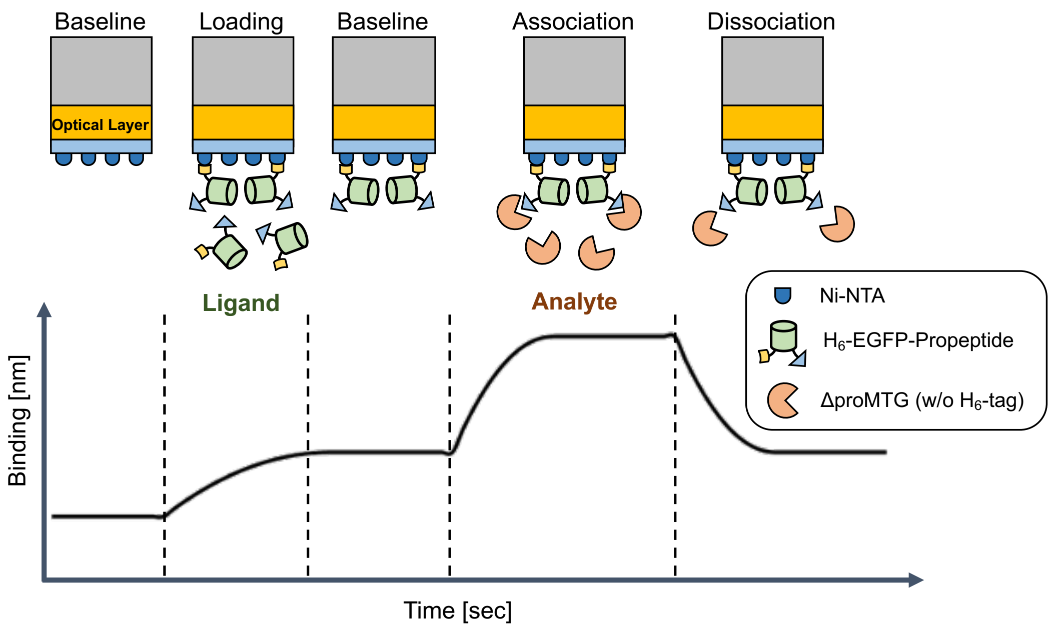


**Fig. S7** Scheme of BLI analysis of the affinity of propeptide with a recombinant matured MTG (ΔproMTG) without His-tag.
